# Near telomere-to-telomere nuclear phased chromosomes of the dikaryotic wheat fungus *Rhizoctonia cerealis*

**DOI:** 10.1101/2022.03.18.484966

**Authors:** Qingdong Zeng, Wenjin Cao, Wei Li, Jianhui Wu, Melania Figueroa, Huiquan Liu, Guowei Qin, Qinhu Wang, Liming Yang, Yan Zhou, Yunxin Yu, Lin Huang, Shengjie Liu, Yuming Luo, Zhiying Mu, Xiang Li, Jiajie Liu, Xiaoting Wang, Changfa Wang, Fengping Yuan, Huaigu Chen, Haibin Xu, Peter N. Dodds, Dejun Han, Zhensheng Kang

## Abstract

*Rhizoctonia cerealis* (*Rce*), which causes sharp eyespot, is one of the most destructive wheat pathogens. However, the genetic and molecular virulence mechanisms of *Rce* have not been elucidated. As a dikaryotic organism, the haplotype phasing of this fungus has not been completed so far. We applied a haplotype phasing algorithm to generate a high-quality near telomere-to-telomere nuclear-phased genome sequence of *Rce* strain R0301. Sixteen pairs of chromosomes were assigned to the A and B genomes with a total size of 83 Mb. Based on a dual-time course RNA-seq, 25308 genes were predicted. Genes for steroid biosynthesis and starch and sucrose metabolism were significantly enriched, together with many genes encoding carbohydrate-active enzymes (CAZymes) and secreted effector proteins, which should be involved in infection of wheat plants. Population genomic analysis of 31 isolates collected in China during the last forty years suggests that this population has not undergone substantial differentiation over time.

**Importance:** The finished genome reference is the basis of revealing pathogens’ biology base. Many efforts have been made to produce the chromosome-scale assembly of fungi. However, the reference of many pathogenic fungi is highly fragmented, which prevents the analysis of genome structure variation, evolution and import pathogenicity genes. Here, we assembly the only chromosome-scale haplotype-phased reference of dikaryotic fungus so far. This assembly achieves the gold standard based on many evaluation software, which indicates that the pipeline developed in this study can be applied to assemble references for other dikaryotic organisms. This work can also promote the research on the globe’s destructive wheat pathogens, sharp eyespot, caused by *R. cerealis*.

---

Basidiomycota fungi with a long-lived dikaryon state play critical ecological roles by degrading plant material and generating significant economic value in the mushroom industry^1^. However, as the dominant causal agents of plant diseases, many fungi of this group also severely threaten agricultural production^2^. As an essential facet of biological research, whole-genome sequences could inspire to solve the most critical and urgent issues on health and agriculture production^3^. Finished genome assemblies present the opportunity to reveal the biology base of pathogens^4^. The genomes of many pathogenic fungi have been reported, while most of these are highly fragmented, preventing the systematic analysis of crucial effector or pathogenicity genes and revealing the information about genome structure variation and evolution^5, 6^. To produce the chromosome-scale assembly of fungi, many attempts have been made, briefly speaking, based on long-read sequencing (Pacbio or Nanopore), combined with genetic mapping, bacterial artificial chromosome or optical mapping^5–9^.

However, these chromosome-scale assembly genomes are mostly reported in monokaryon fungi. For dikaryotic fungi (such as *Puccinia graminis, P. striiformis, P. triticina* and *R. cerealis*), which include species that pose severe threats to global food security, another obstacle to genome assembly has emerged due to the presence of two distinct haploid nuclei in one cell. The collinearity between these divergent haplotypes has often hampered the contiguity of assemblies which have mostly not been able to capture the full sequence information of both haplotypes. A haplotype-phased genome has been generated for only three rust fungi, although these still contain many scaffolding gaps^10, 11^. For *R. cerealis* (anastomosis AG-D group subgroup I (AG-DI)), no reference genome sequence has been reported.

To solve this dilemma on dikaryotic fungi genome assembly, we applied the LAMP assembler algorithm to generate a finished haplotype-phased *R. cerealis* genome sequence based on a combination of Pacbio and nanopore long reads and Illumina short reads. Compared with other available pipelines, our pipeline integrated the contiguity benefit of long reads and the high accuracy of short reads and minimised the occurrence of misassemblies to output a high-quality haplotype phased genome. The single gap in our assembly results from the long tandem repeats (at least 110kbp) associated with a cluster of ribosomal RNA genes, as are usually observed in eukaryotic genomes^12^. Recently, the development of PacBio’s high-fidelity reads produced haplotype-resolved de novo assembly of humans and plants^13–15^. However, due to the short average read length (about 15–20 kb)^13^, this technology may still be hard to fill this gap^16^. On balance, we assembled the only gapless and haplotype-phased reference of a dikaryotic fungus so far, and the pipeline developed in this study can be applied to assemble references for other dikaryotic organisms.

## Results

### Morphological character

*Rce* strain could be cultured on potato dextrose agar (PDA) agar plate, with a white colour of mycelium (Figure 1a), and hyphal cells contained two nuclei each (Figure 1b), confirming that *R. cerealis* is dikaryotic. After inoculation of wheat seedling sheaths and stems, the colour of inoculation sites was converted from white to brown and then black (Figure 1c). In the fields, symptoms on wheat sheaths and stems present semicircular or oval sharp eyespot with grey-brown in the middle and brown around (Figure 1d). The spike of infected wheat was dead and appeared to contain premature ripening whiteheads (Figure 1e).

**Figure 1.**
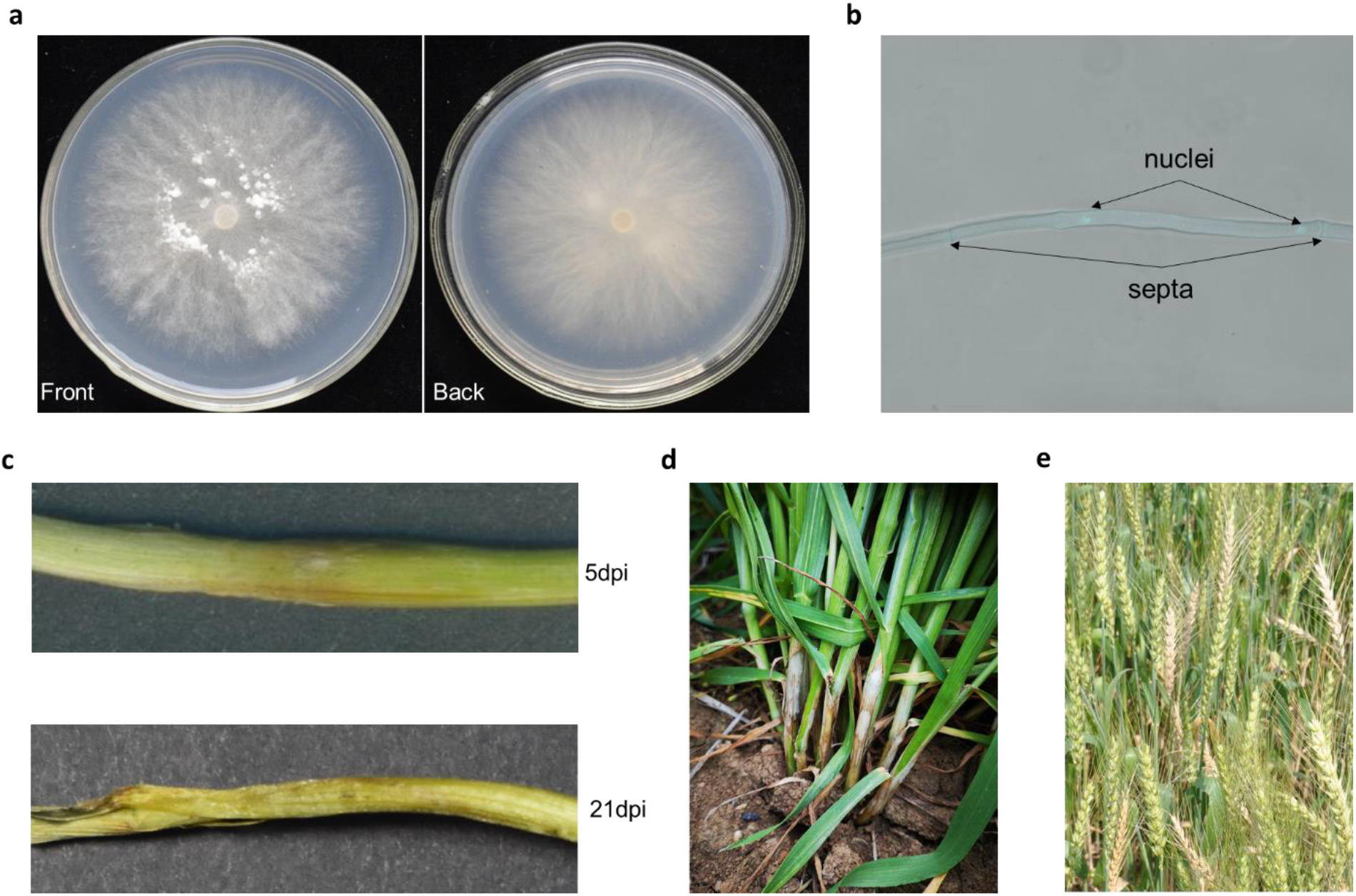
a) The front and backside of R. cerealis hyphal on potato dextrose agar (PDA) agar plat five days post inoculated. b) Fluorescence microscopy of R. cerealis hyphal cells stained with 4',6-diamidino-2-phenylindole (DAPI) demonstrates two nuclei in each cell. c) Symptoms on wheat seedling sheaths and stems at 5 and 21 dpi. d) Field symptoms on wheat sheaths and stems. e) Field symptoms on the wheat spike.

### Genome sequencing and assembly

We generated sequence data from *Rce* genomic DNA using a range of sequencing platform technologies (Supplementary Table 1 at https://figshare.com/s/89e72ddeda3c9e4d8451). A haploid genome size of 40.4 megabase pairs (Mb) of *Rce* was estimated by short reads based on *k*-mer frequency analysis (*k* = 31), thus the genome size of a diploid *Rce* strain was estimated to be nearly 81 Mb in consideration of the heterozygous nature of dikaryotic fungi. The model converged heterozygosity rate is simulated to be 1.19% with two peaks occurring with frequencies of 65× and 132×, respectively (Figure 2a), further confirming the two haploid nuclei of *Rce* differ significantly in genome sequences.

**Figure 2.**
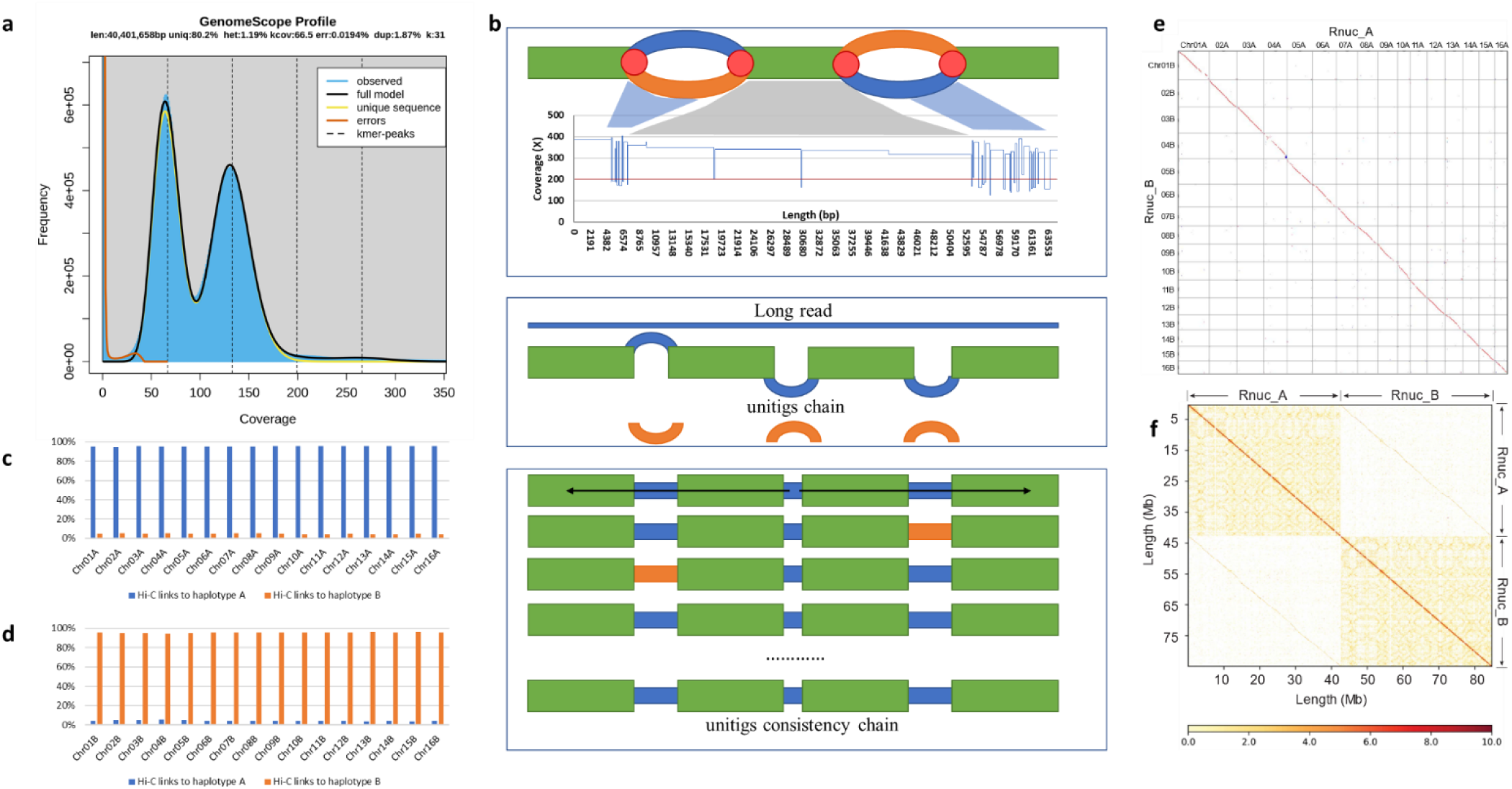
a) GenomeScope k-mer profile plot of the R. cerealis showing the fit of the GenomeScope model (black) to the observed k-mer (k = 31) frequencies (blue). The haploid genome of R. cerealis was estimated to be 40.4 Mb. Two peaks at 65 and 132 were identified. These results indicated that the two haploid nuclei of Rce are highly heterozygous. b) The illustration of the LAMP pipeline. The green bar represents the homozygous region, and the orange or blue bar represents the heterozygous region. The red circle represents the fork point. c) Hi-C read pair links within A genome group. d) Hi-C read pair links within the B genome group. e) Genomic dotplots after 1:1 synteny screen between A and B genome. f) Chromosomal Hi-C contact map data analysis. Inter-chromosomal Hi-C contact map. The intensity of each pixel represents the count of Hi-C links between 100kb windows on chromosomes on a logarithmic scale. Darker red pixels indicate a higher contact probability.

With the same strategy for the primary pool assembly of the short arm of the wheat 2D chromosome (2DS)^17^, we used the LAMP tool to assemble the *Rce* genome (Figure 2b). Briefly, a *de-Bruijn* graph was constructed from short reads (*k*-mer = 99), then all fork points in the graph were broken down to produce the initial unitig set, named NODE. Unitigs from heterozygous regions were identified based on k-mer occurrence frequency, with 150-200× k-mer frequency (Supplementary Figure 1), with these regions representing the two haplotypes assembled separately. On the contrary, unitigs from homozygous regions that were collapsed into a single assembly for the two identical haplotypes, were identified by showing a doubled k-mer frequency of 300-400× and treated as repetitive (Supplementary Figure 1). Unitigs were next aligned to Pacbio long reads, and a chain of unitigs was created for each long read. From these unitig chains, non-repetitive unitigs, or sub-chains with non-repetitive unitigs included, were used as the origin for step-by-step extension to produce haplotype phased genome sequences (Figure 2b).

The *Rce* genome sequence version 0.1 (Rce_V0.1) was produced from Illumina short reads and Pacbio long reads and contained 95 contigs (Supplementary Figure 2). The circular mitochondrial genome was 156,349 bp in length. During the assembly process, extension of contigs was broken off when homozygous regions were encountered with lengths exceeding the available Pacbio long reads from that region (Figure 2b).In the Rce V0.1 assembly, 59 ends from 53 contigs carried a homozygous region with a length of 20 Kb or more (Supplementary Table 2 at https://figshare.com/s/89e72ddeda3c9e4d8451), representing such regions where extension had been halted. This problem was solved by appending Nanopore ultra-long read data in the LAMP pipeline, which reduced the count of contigs to 34, including one derived from the mitochondrion. A total of 54 telomeres with the TTAGG/CCTAA tandem repeat were identified at contig ends (Supplementary Table 3 at https://figshare.com/s/89e72ddeda3c9e4d8451). Nine additional telomeres were identified based on the Pacbio and Nanopore reads aligned to the end of telomere free contigs (Supplemental Note, Supplementary Figure 3). However, due to the low depth of the nine additional telomere sequences supported by Pacbio and Nanopore reads, these telomere sequences were not added to the assembly.

Of the 33 genome contigs, 30 represent entire chromosomes from telomere to telomere (Supplementary Table 3 at https://figshare.com/s/89e72ddeda3c9e4d8451). The three remaining contigs (S9, S18 and S40) contained a telomere at one end but terminated at the other end in long tandem repeats of at least 110kbp (Supplementary Figure 4). These repeats were composed of a cluster of ribosomal RNA genes, as are usually observed in eukaryotic genomes^12^. According to the direction of tandem repeats, S18 was compatible with S40, so these two contigs were joined with a 100bp N linker (Supplemental Note).

No other contig corresponding to the end of S9 was detected, probably because the distance to the telomere is also very short. Thus, our final assembly contained 33 contigs covering 83,241,008 bp (Table 1), representing 16 pairs of homologous chromosomes ranging from 1.5 to 4.0 Mbp and the mitochondrial genome. When the Hi-C read pairs were assessed, the 32 chromosome sequences could be separated into A and B haplotype genomes containing 16 chromosomes each (named Chr01A – Chr16A and Chr01B – Chr16B; Supplementary Figure 5). Chromosome sequences showed a higher proportion of Hi-C read pair links within each haplotype (94.8 - 95.9%) than between haplotypes (4.1 - 5.2%) (Figure 2c,d), consistent with their physical separation in two nuclei. This was further confirmed by the collinearity of the two haploid genomes and whole-genome Hi-C contact map (Figure 2e,f, Supplementary Figure 5). The number of trans read-pair links between contigs S18 and S40 was 39, while only 2 read-pairs connected S18 and S9, which confirmed that S18 and S40 were correctly scaffolded into the same chromosome. This nuclear phased chromosome assembly version was named Rce_V1 and used as the reference of *R. cerealis* strain R0301. For comparison to the LAMP assembly pipeline, we also generated assemblies using several popular algorithms. These all output a very similar genome size from 77.4 to 82.2 Mb (Supplementary Table 4 at https://figshare.com/s/89e72ddeda3c9e4d8451), but with greater fragmentation (higher contig numbers) and containing a number of misjoins and phase swaps between the two nuclear haplotypes (Supplementary Figure 6a to 8a).

**Table 1.** Detail information if the assembly

| Data | Rce_V0.1+nanopore+HiC+PCR |  |  | Pacbio+Illumina |
| --- | --- | --- | --- | --- |
|  | A_genome | B_genome | Rce_V1 | Rce_V0.1 |
| No. of contigs | 16 | 16 | 33 | 95 |
| No. of base | 41,904,800 | 41,336,308 | 83,241,008 | 81,086,359 |
| Max | 3,974,316 | 3,646,437 | 3,974,316 | 2,909,882 |
| N50 | 2,725,423 | 2,408,239 | 2,725,423 | 1,427,415 |
| Median length | 2,537,286 | 2,319,890 | 2,339,717 | 576,440 |
| Average length | 2,619,050 | 2,583,519 | 2,522,455 | 853,541 |
| GC(%) | 48.83 | 48.78 | 48.8 | 48.51 |
| Mito length | - | - | 156,349 | 156,349 |
| TE coverage (%) | 25.22% | 24.06% | 24.66% | 22.01% |
| Quality (QV) | 46.0 | 45.2 | 45.61 | - |
| Completeness | 80.39 | 80.42 | 97.60 | - |

### The assembly of Rce_V1 achieves the gold standard

Four approaches besides the Hi-C contact maps mentioned above were used to evaluate the quality of the Rce_V1 assembly. Firstly, we identified 1,716 (97.3%) complete BUSCOs^18^ (1,701 complete and single-copy BUSCOs) with only 41 missing BUSCOs in the A genome. For the B genome, 1718 (97.4%) complete BUSCOs (1,705 complete and single-copy BUSCOs) and 41 missing BUSCOs were identified (Supplementary Table 4 at https://figshare.com/s/89e72ddeda3c9e4d8451). As non-model fungi, 97% indicates an excellent genome representation. Secondly, all the sequencing reads (Novaseq, PacBio and Nanopore) were mapped against the Rce_V1 assembly, and showed an even read depth across the assembly (Supplementary Figure 9). For novaseq reads, the total mapping rate is 99.41% (properly paired: 98.54%). Thirdly, the correct assembly of long-terminal repeat (LTR) elements was assessed using LTR_retriever^19^, which returned an LTR assembly index (LAI)^20^ value of 23.54, which is classified as the gold quality. Fourthly, based on the spectra copy number plots, the first peak at x=95 and the homozygous content in the second peak at x=190. For nucl A and B (Supplementary Figure 10a, b at https://figshare.com/articles/figure/supplement_figures/19375679), only one haplotype is used, the bubbles in the graph are collapsed, and each heterozygous region is represented once in the assembly. The lost content (the black peak) represents half of the heterozygous content that is lost when bubbles are collapsed. When the whole genome is used, haplotypes are separated by duplicating all the homozygous regions and fully capturing the heterozygous content. According to the KAT^21^ walkthrough, these plots indicated a perfect assembly. At last, based on the k-mers with Merqury^22^, the quality value (QV) score was estimated as 46.0 and 45.2 for haploid A and B genome assembly and and 45.6 for the combined version (Table 1). The K-mer completeness of A, B genome and combined assembly are 80.39, 80.42 and 97.60 (Table 1). The high QV and completeness indicated the assembly of Rce_V1 was excellent performance on quality and completeness.

### Gene prediction and functional annotation

The funannotate pipeline predicted 25,308 genes (26,030 isoforms) (Supplementary Table 5 at https://figshare.com/s/89e72ddeda3c9e4d8451). Among these genes, 1,595 (6.30%) were annotated as carbohydrate activity enzymes (CAZymes) and 2,853 (2,812 representative) candidate secretory effector proteins (CSEP) (Figure 3, Supplementary Table 6 at https://figshare.com/s/89e72ddeda3c9e4d8451). Twenty-four genomic regions were identified as secondary metabolite regions located on 16 chromosomes. Fourteen of these were classified as NRPS/PKS clusters and ten were annotated as terpene-type clusters (Supplementary Figure 11 at https://figshare.com/articles/figure/supplement_figures/19375679, Supplementary Table 7 at https://figshare.com/s/89e72ddeda3c9e4d8451).

**Figure 3.**
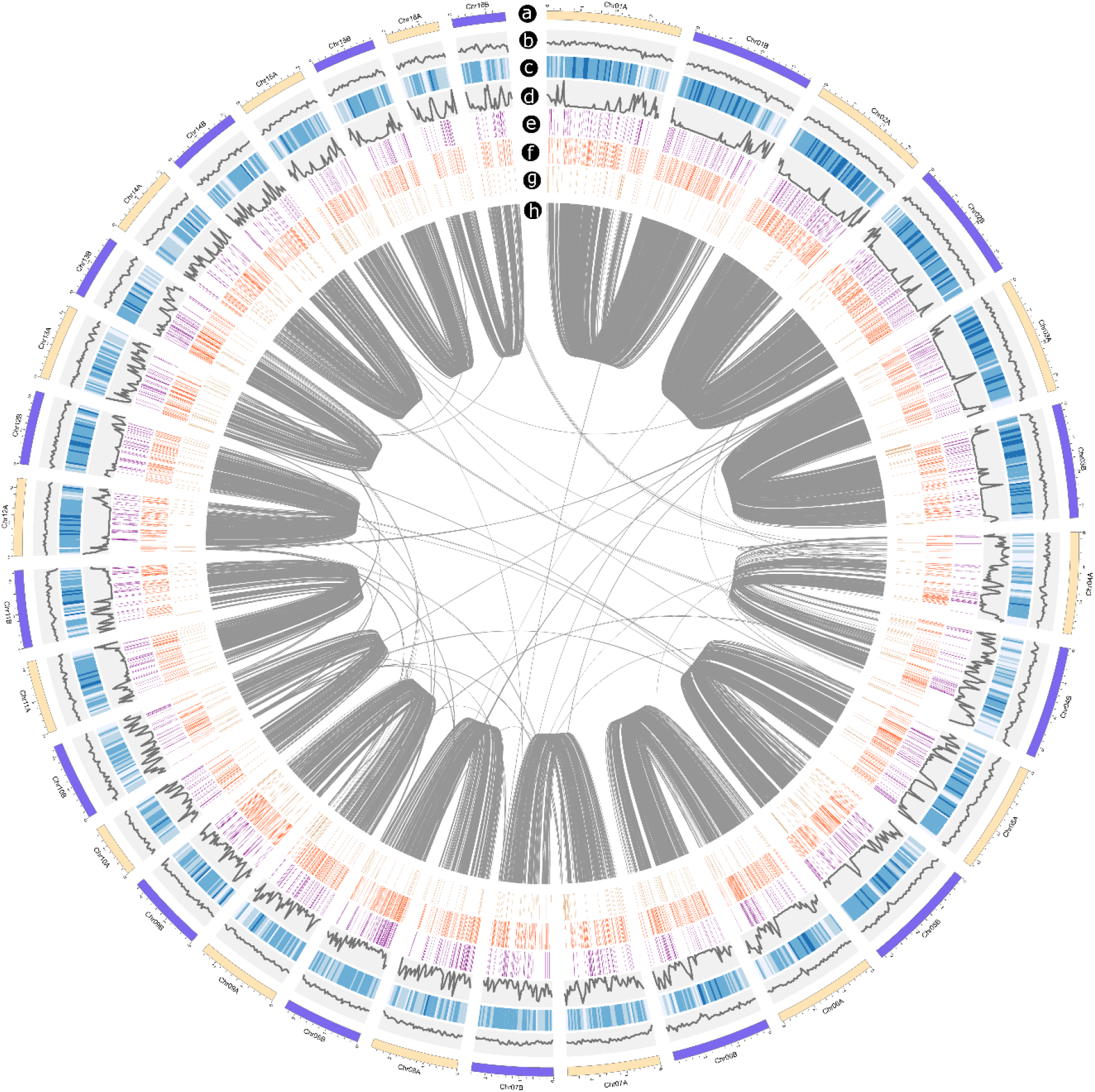
Circos plot of genome features of R. cerealis. a) Karyotype of 32 chromosomes. b) GC content line plotted (0.4-0.6). c) Gene density of 50kb windows. d) Repeat density of 50kb windows. e) Distribution of Cazyme genes. f) Distribution of CSEP genes. g) Distribution of intranuclear specific genes. h) Orthologous gene pairs in the whole genome.

### Haplotype composition and diversity

Repeat sequences make up 25.22% and 24.06% of the A and B genomes respectively. The repeat region distribution is similar between the corresponding A and B chromosomes, but is different among different chromosomes (Figure 3d). The repeat density is higher at the end of most chromosomes, which may cause the incorrect connection among different chromosomes observed for the CANU, FALCON_Unzip and NextDenovo assemblies (Supplementary Figure 6b to 8b). Comparison of the A and B genome haplotypes revealed a total of 718 structural variants affecting about 1.66 Mbp, and 229,003 SNPs, representing 2.75 SNPs/Kb between haplotypes. Compared to *Pgt*, which showed 11–18/Kb SNPs between different haplotypes^10^, the similarity between the A and B haplotypes of *Rce* is much higher. This lower divergence between haplotypes presents a significant for correct phasing and may explain why other assemblers sometimes collapsed two haplotypes into one contig (Supplementary Figure 6c to 8c). Besides the cluster of ribosomal RNA genes (Chr04), we detected 14 other long segment repeats with *k*-mer coverage far above the background genome. These repeats all occurred in subtelomeric regions, with one end of each repeat adjacent to the telomere. Interestingly, all of the 11 such regions in the A genome consisted of an identical 71 Kb sequences, while two regions on Chr01B and Chr11B contained identical 31 Kb sequences and the last region on Chr16B was 14 Kb (Supplementary Table 8 at https://figshare.com/s/89e72ddeda3c9e4d8451, Supplementary Figure 12 at https://figshare.com/articles/figure/supplement_figures/19375679). Since these regions were masked by RepeatMasker and RepeatModeler, the gene was predicted with unmask genome and an additional 277 genes were obtained in these regions (Supplementary Table 9 at https://figshare.com/s/89e72ddeda3c9e4d8451). Twenty-two homolog genes of *Saccharomyces cerevisiae* HST3 are only located on the 11 full-length repeat regions on the A genome. The HST3 gene was reported involved in preventing genome instability^23, 24^, and may be involved in maintaining the stability of the dikaryon system.

The total gene density of both the A and B genomes was 0.3 genes/Kb (Supplementary Table 3 at https://figshare.com/s/89e72ddeda3c9e4d8451). A total of 303 and 347 genes were specific to either the A and B genomes respectively, based on having no hits (-evalue 1e-5) in a reciprocal blastp between A and B genome (Supplementary Table 10 at https://figshare.com/s/89e72ddeda3c9e4d8451). Only a few of these genes are annotated with GO terms, but these were enriched in the biological processes of DNA integration, GPI anchor biosynthetic process, translation, dolichol-linked oligosaccharide (DLO) biosynthetic process and cellular protein modification process. The completely-assembled DLO is the normal N-glycosylation which was reported essential for the activity of *Magnaporthe oryzae*^25^. The function of most haplotype-specific genes was unknown, which may provide clues to reveal the formation and maintenance of the binucleus.

### Transcriptome sequencing and DEG identification

Illumina RNA sequencing (Supplementary Table 11 at https://figshare.com/s/89e72ddeda3c9e4d8451) of *Rce* plants at three time points (at 7,14 and 21 days) growing on PDA medium or in infected plants was conducted. PCA analysis showed that the samples within biological repeats clustered most closely, while the samples collected from infected wheat were also separated from those from PDA, which suggested there may be many genes differentially expressed during the interaction between wheat and *Rce* (Supplementary Figure 13 at https://figshare.com/articles/figure/supplement_figures/19375679). With the GLM (General Linear Models) approach, 10,520 genes were identified as differentially expressed genes (DEG). Separating these genes by their expression trend, 4,457 genes were up-regulated in infected plants versus PDA at the same time point, and 5,327 were down-regulated (Supplementary Table 12 at https://figshare.com/s/89e72ddeda3c9e4d8451). A total of 1,460 genes were commonly up-regulated across all the time points and GO analysis suggested that these were enriched in genes involved in carbohydrate metabolic process, transmembrane transport, sterol biosynthetic process, oxidation-reduction process and cell wall modification. The most enriched pathways were Steroid biosynthesis and Starch and sucrose metabolism (Supplementary Table 13 at https://figshare.com/s/89e72ddeda3c9e4d8451, Supplementary Figure 14 at https://figshare.com/articles/figure/supplement_figures/19375679). Among these 1,460 genes, 298 were annotated as Cazymes, and 413 were annotated as CSEPs, with the proportion of both CAZyme and CSEP significantly higher than in the whole genome (Fisher’s Exact Test, p-value < 0.001). On the other hand, the 1,925 genes down-regulated in all three time points were enriched in protein phosphorylation, oxidation-reduction process, transmembrane transport, methylation, regulation of transcription (DNA-templated), mismatch repair (Supplementary Table 13 at https://figshare.com/s/89e72ddeda3c9e4d8451, Supplementary Figure 14 at https://figshare.com/articles/figure/supplement_figures/19375679). Among these 1,925 genes, there are 127 genes identified as CAZymes and 185 genes were identified as CSEP, which is not significantly different from the proportion in the whole genome (Fisher’s Exact Test, p-value > 0.2). The biological process GO terms of carbohydrate metabolic process and sterol biosynthetic process were significantly and specifically enriched in the commonly up-regulated gene set.

### Population genetics in space and time

Thirty-six isolates of *Rce* were collected from 1984 to 2015 from eight provinces (Supplementary Table 14 at https://figshare.com/s/89e72ddeda3c9e4d8451). After identifying the ITS sequences, 31 isolates from seven provinces in China were sequenced with the Novaseq platform (Supplementary Figure 15a, b at https://figshare.com/articles/figure/supplement_figures/19375679). The A genome of *Rce*_V1 was used as the reference, and the mean mapping rate of these 31 isolates is 93.22% (Supplementary Table 14 at https://figshare.com/s/89e72ddeda3c9e4d8451). With the best practices of the GATK pipeline, 808,864 SNP sites were identified. The filtered VCF (--geno 0.05 --maf 0.05) with 464,887 SNP was used to construct a phylogenetic tree. A reticulated, star-like network output by SplitsTree5 is consistent with extensive recombination among these genotypes due to sexual reproduction (Supplementary Figure 15c at https://figshare.com/articles/figure/supplement_figures/19375679). The PCA analysis divided the population into four groups with two, three, four, and twenty-two isolates (Supplementary Figure 15d at https://figshare.com/articles/figure/supplement_figures/19375679). Based on the tree and PCA, four pairs of likely clonal isolates (very similar genotype) were detected, R1105 and R1404 (three years apart, 300 Km away), R1242 and R1424 (two years apart, 260 Km away), R1301 and R1328 (the same year, 480 Km away), R09130 and R1121 (two years apart, 270 Km away). The clones that appeared between years and field sites indicated that asexual reproduction might also play a significant role in the propagation of *Rce* in the field condition. The linkage disequilibrium (LD) decay at 0.2 was about 3 Kb (Supplementary Figure 15e at https://figshare.com/articles/figure/supplement_figures/19375679) which is consistent with high levels of sexual recombination, although slightly higher than seen in *Zymoseptoria tritici* (less than 1 Kb)^26^, a fungal pathogen that has both asexual and sexual lifestyles, again suggesting an important role for both sexual and asexual reproduction in *Rce*.

### *Rce* specific and significant expansions in energy metabolism and pathogenicity-related genes

Orthofinder was used to identify orthologous gene families between the genomes of *Rce* and *C. theobromae, R. solani* species with default parameters, and the common (at least one gene for all of these seven species) and species-specific families were identified (Figure 4a). There are 292 families (1086 genes) identified as specific to *Rce* A genome. These genes are enriched in the KEGG pathway of ABC transporters, Pentose and glucuronate interconversions, Ribosome biogenesis in eukaryotes, Fructose and mannose metabolism and Pentose phosphate pathway, and Molecular Function GO terms of methyltransferase activity, protein kinase activity, UDP-glycosyltransferase activity, oxidoreductase activity, ATPase-coupled transmembrane transporter activity, iron ion binding, pectate lyase activity, xenobiotic transmembrane transporter activity, oxidoreductase activity, fructose 1,6-bisphosphate 1-phosphatase activity which involved in pathogenicity (Figure 4b). The gene families where one or more species had ≥ 100 gene copies were filtered, and 9143 families were input to cafe (Version 4.2.1)^27^ to generate a species tree (Agabi and CocheC5 as root), which was transformed to the ultrametric tree by r8s. The λ for the whole tree was estimated at 0.000135 to 0.000184. In the *Rce* genome, 1354 families were expansions and 31 of these families were significant, 2062 families were contractions and only one family was significant (Figure 4c). These genes were enriched in MAPK signalling pathway – yeast and Nicotinate and nicotinamide metabolism pathway (Figure 4d). The enriched GO term includes mating-type factor pheromone receptor activity, methyltransferase activity, protein dimerisation activity, protein binding, NAD+ binding (Figure 4d). Overall, *Rce* evolved more nutritional predatory and pathogenic genes to colonise wheat successfully.

**Figure 4.**
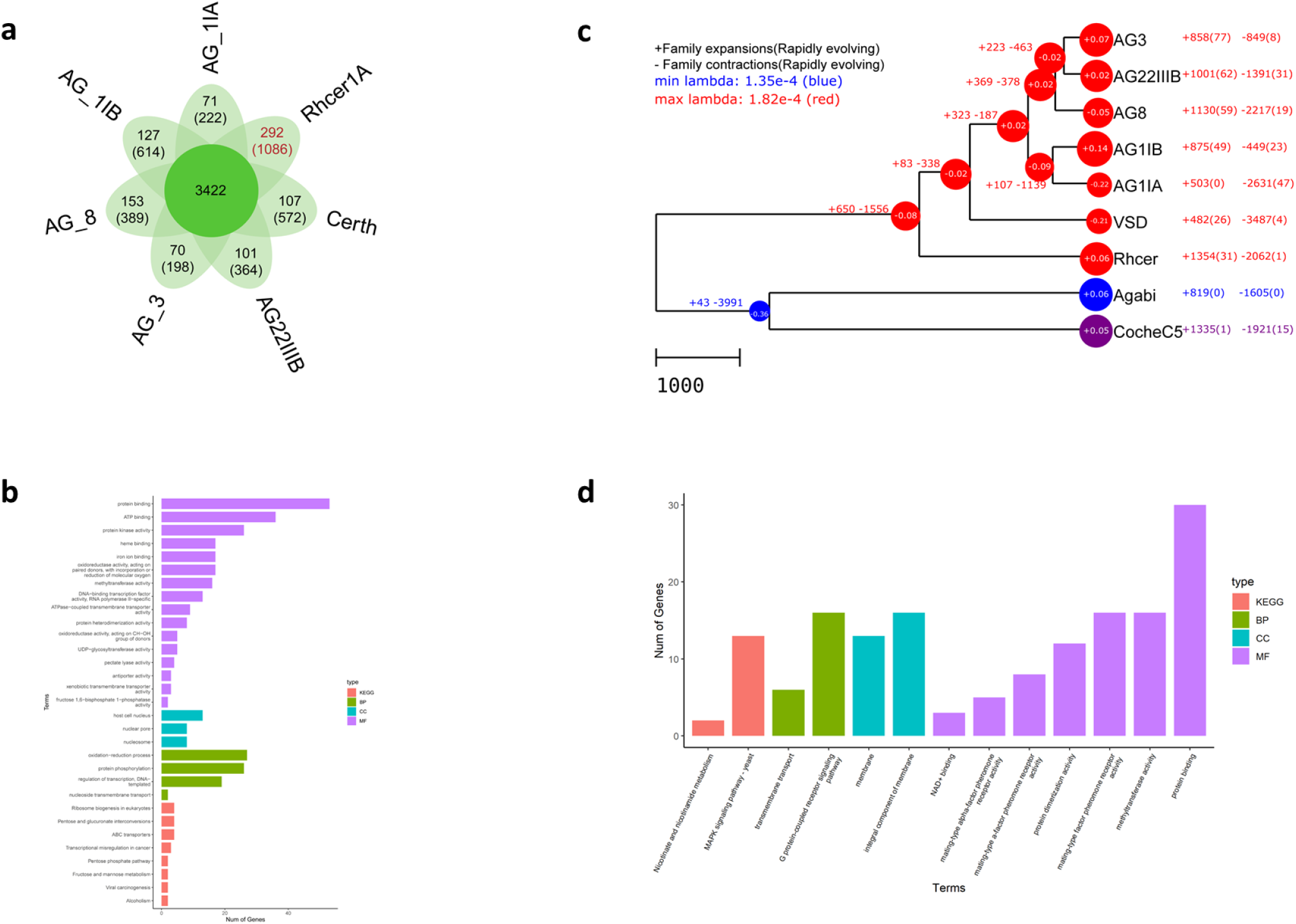
a) Venn diagram of shared and unique orthologue families (genes). b) KEGG and GO enrich the analysis of 1,086 Rhcer A genome-specific genes. c) Evolution of sequenced Rhizoctonia and Ceratobasidium genome. The number in brackets indicated the significant expansions or contraction orthologue family number. d) KEGG and GO enrich analysis of genes in 32 Rhcer A genome significant expansions orthologue family.

### *Rce* sexual development

Though asexual reproduction causes significant yield loss, sexual reproduction plays an important role in recombination^28^, and genetic diversity^29^ and species identity^30^. Sexual fusion between haploid fungal hyphae is controlled by mating type genes^31^, with Basidiomycetes generally containing two mating type loci (*a* and *b*) that encode either a pair of homeodomain transcription factors (HD genes; *b* locus)) or a lipopeptide pheromone and pheromone receptor (PR or P/R genes; *a* locus)^31^. According to DNA binding motifs, the HD proteins were classified into HD1 and HD2 proteins^31^. Based on blast, HMMER and phylogenetic analysis, three allelic pairs of HD genes (two HD1 and one HD2) were identified on Chr15 in a cluster which spans about 12 Kb (Supplementary Figure 16a, b, c at https://figshare.com/articles/figure/supplement_figures/19375679). The protein sequence identity between HD1-2 alleles is 99.80% (RhcerR0301Chr15A02930-T1(HD1-2A) and RhcerR0301Chr15B02680-T1(HD1-2B)), between HD1-1 alleles is 88.19% (RhcerR0301Chr15A02910-T1(HD1-1A) and RhcerR0301Chr15B02660-T1(HD1-1B)) and between HD2 alleles is 82.02% (RhcerR0301Chr15A02920-T1(HD2A) and RhcerR0301Chr15B02670-T1(HD2B)). The protein sequence identity between HD1-1 and HD1-2 was about 66%. All of these genes are expressed with only HD2A, HD2B and HD1-2B significant differentially expressed between cultured and infection samples (Supplementary Table 19 at https://figshare.com/s/89e72ddeda3c9e4d8451). For the pheromone receptor, 34 candidates were identified, 13 of which were removed because less than seven transmembrane domains were identified, leaving 21 genes as the candidant pheromone receptor (STE3). In *Coprinopsis cinerea*, the longest pheromone precursors was 85aa and clustered with the pheromone receptor^31^. To identify pheromone precursors, all of the proteins which length less than 100aa were retrieved and identified the conserved C-terminal motifs CaaX pheromone processing sites. Seven gene RhcerR0301Chr03B10880, RhcerR0301Chr06B06350, RhcerR0301Chr02A05630, RhcerR0301Chr02B05800, RhcerR0301Chr12A06490, RhcerR0301Chr12A06490, RhcerR0301Chr03A10910 were identified as the candidate pheromone precursors. Unfortunately, no identified pheromone receptor and precursors clustered together. An interesting phenomenon is that no clamp connection was observed in the hypha of *Rce*. Based on a previous study, the clamp formation was controlled by the *clp1* gene was induced by the HD protein heterodimer^32–35^. To find some clues about why clampless in *Rce*, the clp1 homologs were identified. However, based on reported clp1 proteins, no conserved regions or motifs were shared^34^. In *Rce*, three gene pairs were identified as *clp1* homologs: RhcerR0301Chr03B08840-T1, RhcerR0301Chr03A08870-T1, RhcerR0301Chr03B01250-T1, RhcerR0301Chr03A01330-T1, RhcerR0301Chr03B01300-T1 and RhcerR0301Chr03A01380-T1 (Supplementary Figure 16d at https://figshare.com/articles/figure/supplement_figures/19375679). Based on transcripts Per Million (TPM) data, all of these genes are significantly differentially expressed with and without wheat induced (Supplementary Table 19 at https://figshare.com/s/89e72ddeda3c9e4d8451).

## Discussion

So far, chromosome-level genome assembly has been available in many haploid and inbred species^5–9^, while phasing of non-inbred or rearranged heterozygous genomes has posed great challenges^36^. Here we report a chromosome-scale haplotype phased *Rce* genome based on nearly all sequencing platforms except bionano and sanger, which were high-cost input. Our assembly has an essential improvement compared to the published eight *R. solani* and one *Ceratobasidium theobromae* genome (Supplementary Table 16 at https://figshare.com/s/89e72ddeda3c9e4d8451). The high collinearity between the two haplotypes of the dikaryon and the high repeat density in the chromosome impeded the correct assembly with widely used assemblers. In contrast, the LAMP assembler integrated the benefit of long read length of Pacbio and Nanopore data and the high accuracy of Illumina short reads and minimised the occurrence of mis-assemblies by manual auxiliary judgement in real-time and output a high-quality haplotype phased genome. We also provide the first report of long segment repeats that show nucelus-specific distribution in this genome. On the one hand, this work provides the high-quality reference of *Rce* and the clue of the dikaryon system stability, which will promote the research on genomics, functioning, evolution, and disease control for this organism. On the other hand, this study provides a new method and strategy for complex dikaryon fungi genome assembly.

The nuclear phased chromosome-scale reference also provides the blueprint to assess variation between the two karyons. The SNP density between the A and B genome was much smaller than the SNP variation among isolates (229,003 vs 808,864), which indicated that free recombination occurred. Although no sexual stage has been found in the field or the laboratory^37^, it is hard to explain without sexual reproduction why existed high genetic and genotypic diversity in the field both emerged in this and previous studies^38, 39^. Paired clones collected in a different year and from distant fields indicated that asexual reproduction likely occurs and might play an important role in the disease. Based on the whole genome resequencing data, this study found that 31 isolates collected in China spanning nearly 40 years from seven provinces had no evident clustering. The evolutionary tree is not separated by year or province. These results suggest there is significant gene flow among different provinces, which may result from the infected seed or harvester spread^38^. No regular clustering according to space and time also indicated the population as a whole has not undergone substantial differentiation over time. This may be due to limited selection imposed by host resistance and lack of introgression of new *Rce* isolates from other global populations. There are two possible reasons, firstly, the lack of resistant cultivars in wheat production with only chemical control used over this period may mean there has been little selection to adapt to host resistance genes^37, 38^. Secondly, the 40-year time span may be too short to detect selection. Overall, the *Rce* may have asexual predominates and sexual mixed reproduction strategies to maintain the genetic diversity and the selective pressure was too low to promote the formation of isolate clusters in specific time and space.

The vascular plant cell wall is the first barrier against fungal pathogens and is composed of polysaccharides (cellulose, hemicellulose, and lignin), phenolic compounds and proteins^1, 40^. Many research works reported that the composition and structures of the cell wall contributes to resist pathogen invasion^41–44^. To colonise successfully, necrotrophic fungi produce different CAZymes like polygalacturonases, hemicellulases, cellulases, acetylesterases and pectin methylesterases^42^. These enzymes degrade cell wall polymers to facilitate infection and supply nutrition for fungal growth^45^. Genes significantly Up-regulated in all the interaction samples compared to the samples collected from PDA were enriched in carbohydrate metabolic process, transmembrane transport, sterol biosynthetic process, oxidation-reduction process and cell wall modification biological process. The genes involved in nutritional predatory and pathogenicity enriched in *Rce* specific and significate expansions gene set. These results indicate *Rce* probable produce CAZymes to degrade wheat cell wall for colonisation, and at the same time, the generative saccharides could be transported into fungus and used as a carbon source. This result provides the theoretical basis for manipulating host sugar metabolism and transport to control this disease.

## Materials and Methods

Please see the online version.

## Acknowledgements

The authors thank Hua Zhao (NWAFU) for assisting on microscopy and Guoliang Pei (NWAFU) for assisting on cluster management. This work was supported by grants from the National Natural Science Foundation of China (31600444, 31620103913), by grants from the Opening Foundation of the Jiangsu Key Laboratory for Eco-Agricultural Biotechnology Around Hongze Lake (HZHLAB2001), by grants from the Opening Foundation of the State Key Laboratory of Crop Stress Biology for Arid Areas (CSBAAKF2018001), by grants from Hong-Yuan Biotech Co., Ltd (Nanjing, China), by grants from BioCAM Biotech Co., Ltd (Nanjing, China), by grants from National ‘111 plan’ (BP0719026).

## Author Contributions

The project was principally investigated by HX, DH, ZK, PND and HC. Sequences were assembled by HX and QZ. Bioinformatics analyses were performed by QZ, PND, MF, JW, WL and LY. Experiments for validation were performed by WC, SL, FY, YZ, YY, LH, XL, JL, XW, CW, GQ, YL and ZM. The paper was written by QZ, HX, MF, PND and was revised by DH, ZK, JW, HL and WL. HL guided sexual development analysis. QW contributed the evolution analysis. All authors read and commented on the manuscript.

## Competing Interests

The authors declare no competing interests.

## Data availability

All sequence data, assemblies generated in this study are available in NCBI under BioProject PRJNA717151. Assemblies and other data have been deposited at figshare (https://figshare.com/) with DOI numbers of 10.6084/m9.figshare.14256686 (*Rce* unitigs *private link:* https://figshare.com/s/0c06a42ebe1ec49b1aaf), 10.6084/m9.figshare.14256674 (*Rce* Lamp assembly *private link:* https://figshare.com/s/d3533018091c33cb77f2}, 10.6084/m9.figshare.14256644 (*Rce* Canu assembly *private link:* https://figshare.com/s/8d6478a2aa52d69a1386), 10.6084/m9.figshare.14256662 (*Rce* Falcon_unzip assembly *private link:* https://figshare.com/s/82e5530294e7d63289c3) 10.6084/m9.figshare.14256656 (Falcon_phase assembly *private link:* https://figshare.com/s/3006d34e80e5ecb25e61) 10.6084/m9.figshare.14256677 (*Rce* NextDenovo assembly *private link:* https://figshare.com/s/34b885730f704b289b10) 10.6084/m9.figshare.14256680 (*Rce* Supernova assembly *private link:* https://figshare.com/s/478762f9d544bb228752)

## Supplementary Table

Supplementary Table 1 Summary of genomic sequencing data of R0301. a) Summary of Novaseq genomic sequencing data of R0301. b) Summary of Pacbio genomic sequencing data of R0301. c) Summary of Hi-C genomic sequencing clean data of R0301. d) Summary of 10x genomics sequencing clean data of R0301. e) Summary of Nanopore genomics sequencing clean data of R0301. Post at https://figshare.com/s/89e72ddeda3c9e4d8451

Supplementary Table 2 The length of homozygous regions of *Rce*_V0.1 contigs end which larger than 20 kb. Post at https://figshare.com/s/89e72ddeda3c9e4d8451

Supplementary Table 3 The detailed information of each chromosome. Post at https://figshare.com/s/89e72ddeda3c9e4d8451

Supplementary Table 4 The detailed assembly information of different pipeline. Post at https://figshare.com/s/89e72ddeda3c9e4d8451

Supplementary Table 5 Gene prediction and function annotation of *Rce* Post at https://figshare.com/s/89e72ddeda3c9e4d8451

Supplementary Table 6 Candidate secretory effector proteins of *Rce*. Post at https://figshare.com/s/89e72ddeda3c9e4d8451

Supplementary Table 7 Identified secondary metabolite regions and the core biosynthetic genes. Post at https://figshare.com/s/89e72ddeda3c9e4d8451

Supplementary Table 8 The information of 14 long segment repeats. Post at https://figshare.com/s/89e72ddeda3c9e4d8451

Supplementary Table 9 Gene and function annotation in the long repeat regions. Post at https://figshare.com/s/89e72ddeda3c9e4d8451

Supplementary Table 10 Haplotype specific genes and annotation. Post at https://figshare.com/s/89e72ddeda3c9e4d8451

Supplementary Table 11 Wheat and *R. cerealis* interaction dual Illumina RNA seq and Pacbio iso seq. Post at https://figshare.com/s/89e72ddeda3c9e4d8451

Supplementary Table 12 Differentially expressed gene identified between wheat and *Rce* interaction. Post at https://figshare.com/s/89e72ddeda3c9e4d8451

Supplementary Table 13 GO and KEGG enrich analysis result of DEG between wheat and *Rce* interaction. Post at https://figshare.com/s/89e72ddeda3c9e4d8451

Supplementary Table 14 Summary of resequencing data of 31 isolates. Post at https://figshare.com/s/89e72ddeda3c9e4d8451

Supplementary Table 15 The genome information of *Rhizoctonia solani*. Post at https://figshare.com/s/89e72ddeda3c9e4d8451

Supplementary Table 16 Compare to eight published *R. solani* and one *Ceratobasidium theobromae* genome. Post at https://figshare.com/s/89e72ddeda3c9e4d8451

Supplementary Table 17 Different local alternative splicing event types generated by SUPPA based on Iso-seq data. Post at https://figshare.com/s/89e72ddeda3c9e4d8451

Supplementary Table 18 The detailed information about the genes involved in sexual development. Post at https://figshare.com/s/89e72ddeda3c9e4d8451

Supplementary Table 19 The expression of the genes involved in sexual development in *Rce*. Post at https://figshare.com/s/89e72ddeda3c9e4d8451

## Supplementary Figure

Supplementary Figure 1 Flowchart of the workflow used to estimate *k*-mer coverages for heterozygous and homozygous genomic regions in the diploid *R. cerealis* genome. Each Illumina short read was end-trimmed to 125 bp in length, then disassembled to produce 27 99-bp *k*-mers. A total of 16 billion *k*-mers were produced from 593 million reads. For the 81 Mb diploid *R. cerealis* genome, one-fold *k*-mer coverage would be approximately 81 million *k*-mers. Thus, the disassembled *k*-mers would correspond to about 198× average coverage depth for the *R. cerealis* genome. Adjusting this estimate to account for sources of contamination, including from mitochondrion, endophytic microorganisms, etc., coverage is an estimated 150-200× for heterozygous genomic regions and 300-400× for homozygous genomic regions.

Supplementary Figure 2 Flowchart of the process used to assemble genome sequences of the dikaryotic *R. cerealis* strain R0301 by utilising the Lamp algorithm based on the WGS data.

Supplementary Figure 3 The Nanopore and Pacbio reads failed assembled telomere in the nine chromosome ends.

Supplementary Figure 4 The tandem repeats at the end of S18 and S40 contigs. a) The dotplot of S40 vs S40 contig. b) The dotplot of Nanopore reads 7438396d-f174-43a6-8bf4-1876afbb1abc which is local at the distal telomere end of S40. c) The dotplot of S18 vs S18 contig. d) The dotplot of nanopore reads 3d603e88-7e3a-45ca-9ae8-2a1a6014c347 which local at the distal telomere end of S18.

Supplementary Figure 5 Intra-chromosomal contact maps of each chromosome. The intensity of each pixel represents the count of Hi-C links between 20kb windows on each chromosome on a logarithmic scale. Darker red pixels indicate a greater contact probability.

Supplementary Figure 6 Illustration of mis-assemblies in the Canu assembly of the *R. cerealis* genome: a) Global comparison of the Canu assembly to our chromosome assembly (Rce_v1), ordered by chromosome. The grey and coloured rectangles mark alignments corresponding to each chromosome pair. The Canu assembly consists of 520 contigs with a total length of 81.0 Mb. Canu assembly contigs corresponding to each chromosome pair can be seen on y-axis labels and generally consist of a few long contigs and many short contigs for each chromosome pair. Alignments marked with orange or green rectangles were chosen as the examples to enlarge as follows to inspect mis-assemblies in the Canu assembly further. Contigs less than 100 Kb in the Canu assembly was removed. b) Portions of chromosomes 03A and 08B were erroneously joined to produce the tig00001353 contig in the Canu assembly (left graph). Blue rectangles indicate the junction position of this mis-assembly in dot plots and a bold red line in the genome browser view. The erroneous nature of this mis-assembly is supported by the contact break shown in the Hi-C contact map for this contig (middle graph) and by the drop in coverage for Illumina PE reads, PacBio CLR reads and Nanopore ultra-long reads (right graph). c) Haplotypes were collapsed in portions of the Canu assembly corresponding to chromosome pair No. 01, leading to three gaps with lengths over 200 kb in the alignment to chromosomes 01A and 01B (Chr01A:2,627-2,829 kb, Chr01B:914-1,236 kb and Chr01B:1,495-1,712 kb). Primary alignments showing the highest identity were concentrated in the region unmarked by grey triangles. Blue rectangles mark the position of the three gaps in the Rce_v1 assembly.

Supplementary Figure 7 Illustration of mis-assemblies in the Falcon_unzip assembly of the *R. cerealis* genome with colour schemes identical to those in Supplementary Figure 6: a) Global comparison of the Falcon_unzip assembly to our chromosome assembly (Rce_v1). The Falcon_unzip assembly consists of 280 contigs with a total length of 82.2 Mb. Similar to the Canu assembly, contig groups corresponding to each chromosome pair generally consist of a few long contigs and many shorter contigs. Contigs less than 100 Kb in the Falcon_unzip assembly was removed. b) The chromosomes 03B and 01A were erroneously joined to produce the 000012F|arrow contig in the Falcon_unzip assembly (left graph). The erroneous nature of this mis-assembly is supported both by the contact break shown in the Hi-C contact map for this contig (middle graph) and by the drop in PacBio CLR read coverage at the junction (right graph). c) Portions of haplotypes were collapsed or misplaced in chromosome pair No. 01 in the Falcon_unzip assembly, leading to three gaps of over 100 kb (Chr01A:3,649-3,784 kb, Chr01B:747-1,282 kb and Chr01B:1,354-1,487 kb). The only gap that overlaps with the Canu assembly gap presented in Figure s21c is the gap in the middle position.

Supplementary Figure 8 Illustration of mis-assemblies in the NextDenovo assembly of *R. cerealis* genome with colour schemes identical to those shown in Supplementary Figure 6: a) Global comparison of the NextDenovo assembly to our chromosome assembly (Rce_v1). The NextDenovo assembly consists of 61 contigs, with a total length of 77.4 Mb, notably smaller than Rce_v1 (83.2 Mb). The mitochondrial genome was absent from the NextDenovo assembly. The red arrow marks a mis-assembly in the ctg000037_np512 contig, which features an errant joining of chromosomes 09A and 09B in an end-to-end manner. b) The chromosomes 04B and 08A were erroneously joined to produce the ctg000001_np512 contig in the NextDenovo assembly (left graph). The erroneous nature of this mis-assembly is supported both by the contact break shown in the Hi-C contact map for this contig (middle graph) and by the drop in coverage at the junction position with Illumina PE reads, PacBio CLR reads and Nanopore ultra-long reads (right graph). c) Internal portions of haplotypes were collapsed in chromosome pair No. 01, leading to the ctg000018_np512 contig representing two internal portions on chromosomes 01A and 01B, with the length of 2.0 Mb for each.

Supplementary Figure 9 The visualisation of mapped depth all the used reads against the Rce_V1 assembly with 1 Kb window. a) A genome used as the reference. b) B genome used as the reference. Top, Illumina Novaseq PE reads. Middle, Nanopore ultra-long reads. Bottom, PacBio CLR reads.

Supplementary Figure 10 K-mer spectra copy number plot of Rce_V1 assembly based on the Illumina Novaseq PE reads. The first peak represents the heterozygous content at x=90 and the homozygous content at the second peak at x=180. The lost content (the black peak) represents half of the heterozygous content lost when bubbles collapse. a) Only the nucl A genome haplotype was used. b) Only the nucl B genome haplotype was used. (c) The whole genome was used as the reference. Post at https://figshare.com/articles/figure/supplement_figures/19375679

Supplementary Figure 11 Genome-wide secondary metabolite regions identified in *Rce*. Post at https://figshare.com/articles/figure/supplement_figures/19375679Supplementary Figure 12 The alignment of the 14 long repeat regions. Post at https://figshare.com/articles/figure/supplement_figures/19375679

Supplementary Figure 13 MDS plot of 18 samples based on logFC with BCV method. a) Wheat is grown with and without *Rce* infected. b) MDS plot of all samples. Post at https://figshare.com/articles/figure/supplement_figures/19375679

Supplementary Figure 14 Veen and enrich the analysis of DEG between wheat and *Rce* interaction. a) up-grade genes and annotation. b) down-grade genes and annotation. Post at https://figshare.com/articles/figure/supplement_figures/19375679

Supplementary Figure 15 31 sequenced isolates information. a) The sample location of 31 isolates from China. b) The phylogenetic tree is based on the ITS sequences. c) The reticulated, star-like network based on the genome-wide SNP data. d) The PCA analysis of the population. e) The linkage disequilibrium (LD) decay. Post at https://figshare.com/articles/figure/supplement_figures/19375679

Supplementary Figure 16 Homeodomain genes identified. a) The phylogenetic tree of HD1 genes. b) The phylogenetic tree of HD2 genes. c) The location and structure of HD genes. d) The phylogenetic tree of *clp1* genes. Post at https://figshare.com/articles/figure/supplement_figures/19375679

## References

1. Morin, E. et al. Genome sequence of the button mushroom *Agaricus bisporus* reveals mechanisms governing adaptation to a humic-rich ecological niche. Proceedings of the National Academy of Sciences of the United States of America 109, 17501–17506 (2012).

2. Sperschneider, J. et al. Advances and challenges in computational prediction of effectors from plant pathogenic fungi. PLoS Pathog 11, e1004806 (2015).

3. Sharma, K.K. Fungal genome sequencing: basic biology to biotechnology. Crit. Rev. Biotechnol. 36, 743–U227 (2016).

4. Koren, S. et al. Canu: scalable and accurate long-read assembly via adaptive k-mer weighting and repeat separation. Genome Res. 27, 722–736 (2017).

5. Muller, M.C. et al. A chromosome-scale genome assembly reveals a highly dynamic effector repertoire of wheat powdery mildew. New Phytol. 221, 2176–2189 (2019).

6. Faino, L. et al. Single-Molecule Real-Time Sequencing Combined with Optical Mapping Yields Completely Finished Fungal Genome. MBio 6 (2015).

7. Nierman, W.C. et al. Genomic sequence of the pathogenic and allergenic filamentous fungus *Aspergillus fumigatus*. Nature 438, 1151–1156 (2005).

8. Torriani, S.F., Goodwin, S.B., Kema, G.H., Pangilinan, J.L. & McDonald, B.A. Intraspecific comparison and annotation of two complete mitochondrial genome sequences from the plant pathogenic fungus *Mycosphaerella graminicola*. Fungal Genet. Biol. 45, 628–637 (2008).

9. Goodwin, S.B. et al. Finished Genome of the Fungal Wheat Pathogen *Mycosphaerella graminicola* Reveals Dispensome Structure, Chromosome Plasticity, and Stealth Pathogenesis. PLoS Genet. 7 (2011).

10. Li, F. et al. Emergence of the Ug99 lineage of the wheat stem rust pathogen through somatic hybridisation. Nature Communications 10 (2019).

11. Duan, H. et al. Identification and correction of phase switches with Hi-C data in the Nanopore and HiFi chromosome-scale assemblies of the dikaryotic leaf rust fungus *Puccinia triticina*. bioRxiv, 2021.2004.2028.441890 (2021).

12. Rabanal, F.A. et al. Epistatic and allelic interactions control expression of ribosomal RNA gene clusters in *Arabidopsis thaliana*. Genome Biol 18, 75 (2017).

13. Wenger, A.M. et al. Accurate circular consensus long-read sequencing improves variant detection and assembly of a human genome. Nat. Biotechnol. 37, 1155–1162 (2019).

14. Cheng, H., Concepcion, G.T., Feng, X., Zhang, H. & Li, H. Haplotype-resolved de novo assembly using phased assembly graphs with hifiasm. Nat. Methods (2021).

15. Nurk, S. et al. HiCanu: accurate assembly of segmental duplications, satellites, and allelic variants from high-fidelity long reads. Genome Res. 30, 1291–1305 (2020).

16. Garg, S. Computational methods for chromosome-scale haplotype reconstruction. Genome Biol 22, 101 (2021).

17. Zeng, Q. et al. A novel high-accuracy genome assembly method utilizing a high-throughput workflow. bioRxiv, 2020.2011.2026.400507 (2020).

18. Waterhouse, R.M. et al. BUSCO applications from quality assessments to gene prediction and phylogenomics. Mol. Biol. Evol. (2017).

19. Ou, S. & Jiang, N. LTR_retriever: A Highly Accurate and Sensitive Program for Identification of Long Terminal Repeat Retrotransposons. Plant Physiol. 176, 1410–1422 (2018).

20. Ou, S., Chen, J. & Jiang, N. Assessing genome assembly quality using the LTR Assembly Index (LAI). Nucleic Acids Res. 46, e126 (2018).

21. Mapleson, D., Garcia Accinelli, G., Kettleborough, G., Wright, J. & Clavijo, B.J. KAT: a K-mer analysis toolkit to quality control NGS datasets and genome assemblies. Bioinformatics 33, 574–576 (2017).

22. Rhie, A., Walenz, B.P., Koren, S. & Phillippy, A.M. Merqury: reference-free quality, completeness, and phasing assessment for genome assemblies. Genome Biol 21, 245 (2020).

23. Hachinohe, M., Hanaoka, F. & Masumoto, H. Hst3 and Hst4 histone deacetylases regulate replicative lifespan by preventing genome instability in *Saccharomyces cerevisiae*. Genes Cells 16, 467–477 (2011).

24. Feldman, J.L. & Peterson, C.L. Yeast Sirtuin Family Members Maintain Transcription Homeostasis to Ensure Genome Stability. Cell Reports 27, 2978–2989.e2975 (2019).

25. Chen, X.-L. et al. *N*-Glycosylation of Effector Proteins by an α-1,3-Mannosyltransferase Is Required for the Rice Blast Fungus to Evade Host Innate Immunity. The Plant Cell 26, 1360 (2014).

26. Singh, N.K., Karisto, P. & Croll, D. Population-level deep sequencing reveals the interplay of clonal and sexual reproduction in the fungal wheat pathogen *Zymoseptoria tritici*. Microb. Genomics 7 (2021).

27. Han, M.V., Thomas, G.W., Lugo-Martinez, J. & Hahn, M.W. Estimating gene gain and loss rates in the presence of error in genome assembly and annotation using CAFE 3. Mol. Biol. Evol. 30, 1987–1997 (2013).

28. Rice, W.R. Experimental tests of the adaptive significance of sexual recombination. Nat. Rev. Genet. 3, 241–251 (2002).

29. Agrawal, A.F. Evolution of sex: Why do organisms shuffle their genotypes? Curr. Biol. 16, R696–R704 (2006).

30. Heng, H.H. Elimination of altered karyotypes by sexual reproduction preserves species identity. Genome 50, 517–524 (2007).

31. Kues, U. From two to many: Multiple mating types in Basidiomycetes. Fungal Biol Rev 29, 126–166 (2015).

32. Inada, K., Morimoto, Y., Arima, T., Murata, Y. & Kamada, T. The *clp1* gene of the mushroom *Coprinus cinereus* is essential for A-regulated sexual development. Genetics 157, 133–140 (2001).

33. Kamada, T. Molecular genetics of sexual development in the mushroom *Coprinus cinereus*. Bioessays 24, 449–459 (2002).

34. Scherer, M., Heimel, K., Starke, V. & Kamper, J. The Clp1 protein is required for clamp formation and pathogenic development of *Ustilago maydis*. Plant Cell 18, 2388–2401 (2006).

35. Ekena, J.L., Stanton, B.C., Schiebe-Owens, J.A. & Hull, C.M. Sexual development in *Cryptococcus neoformans* requires *CLP1*, a target of the homeodomain transcription factors Sxi1alpha and Sxi2a. Eukaryot. Cell 7, 49–57 (2008).

36. Chin, C.S. et al. Phased diploid genome assembly with single-molecule real-time sequencing. Nat. Methods 13, 1050–1054 (2016).

37. Hamada, M.S., Yin, Y., Chen, H. & Ma, Z. The escalating threat of *Rhizoctonia cerealis*, the causal agent of sharp eyespot in wheat. Pest Manag Sci 67, 1411–1419 (2011).

38. Li, W., Guo, Y.P., Zhang, A.X. & Chen, H.G. Genetic Structure, of Populations of the Wheat Sharp Eyespot Pathogen *Rhizoctonia cerealis* Anastomosis Group D Subgroup I in China. Phytopathology 107, 224–230 (2017).

39. Liu, J. & Mundt, C.C. Genetic structure and population diversity in the wheat sharp eyespot pathogen *Rhizoctonia cerealis* in the Willamette Valley, Oregon, USA. Plant Pathol. 69, 101–111 (2020).

40. Yokoyama, R. & Nishitani, K. Genomic basis for cell-wall diversity in plants. A comparative approach to gene families in rice and *Arabidopsis*. Plant Cell Physiol. 45, 1111–1121 (2004).

41. Pogorelko, G., Lionetti, V., Bellincampi, D. & Zabotina, O. Cell wall integrity: targeted post-synthetic modifications to reveal its role in plant growth and defense against pathogens. Plant Signal Behav 8 (2013).

42. Bellincampi, D., Cervone, F. & Lionetti, V. Plant cell wall dynamics and wall-related susceptibility in plant-pathogen interactions. Front Plant Sci 5, 228 (2014).

43. Lionetti, V. & Metraux, J.P. Plant cell wall in pathogenesis, parasitism and symbiosis. Front Plant Sci 5, 612 (2014).

44. Basińska-Barczak, A., Błaszczyk, L. & Szentner, K. Plant Cell Wall Changes in Common Wheat Roots as a Result of Their Interaction with Beneficial Fungi of *Trichoderma*. Cells 9, 2319 (2020).

45. Blanco-Ulate, B. et al. Genome-wide transcriptional profiling of *Botrytis cinerea* genes targeting plant cell walls during infections of different hosts. Front Plant Sci 5, 435 (2014).

